# Early Archean origin of heterodimeric Photosystem I

**DOI:** 10.1101/225888

**Authors:** Tanai Cardona

**Affiliations:** Department of Life Sciences, Imperial College London, London, UK

## Abstract

When and how oxygenic photosynthesis originated remains controversial. Wide uncertainties exist for the earliest detection of biogenic oxygen in the geochemical record or the origin of water oxidation in ancestral lineages of the phylum Cyanobacteria. A unique trait of oxygenic photosynthesis is that the process uses a Type I reaction centre with a heterodimeric core, also known as Photosystem I, made of two distinct but homologous subunits, PsaA and PsaB. In contrast, all other known Type I reaction centres in anoxygenic phototrophs have a homodimeric core. A compelling hypothesis for the evolution of a heterodimeric Type I reaction centre is that the gene duplication that allowed the divergence of PsaA and PsaB was an adaptation to incorporate photoprotective mechanisms against the formation of reactive oxygen species, therefore occurring after the origin of water oxidation to oxygen. Here I show, using sequence comparisons and Bayesian relaxed molecular clocks that this gene duplication event may have occurred in the early Archean more than 3.4 billion years ago, long before the most recent common ancestor of crown group Cyanobacteria and the Great Oxidation Event. If the origin of water oxidation predated this gene duplication event, then that would place primordial forms of oxygenic photosynthesis at a very early stage in the evolutionary history of life.

## Introduction

When and how oxygenic photosynthesis originated remains a highly debated subject with dates ranging from 3.8 billion years (Ga) (1, 2) to shortly before 2.4 Ga (3, 4), the onset of the Great Oxidation Event (GOE). Extensive geochemical data suggests that some oxygen was being produced from at least about 3.0 Ga (5-7). Yet, it is well accepted that anoxygenic photosynthesis had already evolved 3.5 Ga ago and may predate 3.8 Ga (8-10). If so, there could have been a gap of 600 million years (Ma) to more than 1.0 Ga between the origin of anoxygenic photosynthesis and the emergence of oxygenic photosynthesis. However, I have previously pointed out that the molecular evolution of photochemichal reaction centres is inconsistent with such a vast gap (11-13). Instead, I have noted that the evolution of all reaction centre proteins, including those used in oxygenic photosynthesis, diversified early, rapidly, and in parallel (14). Consistent with this, Cardona et al. (13) demonstrated that the divergence of the anoxygenic Type II reaction centres from water-oxidising Photosystem II (PSII) occurred soon after the emergence of photosynthesis as the result of one reaction centre gaining or losing the structural domains required to oxidise water to oxygen, still at a homodimeric stage. Cardona et al. (13) also showed that one of the most ancient diversification events in the evolution of photosynthesis, after the divergence of Type I and Type II reaction centres, was the specific gene duplication of the ancestral water-splitting homodimeric core that permitted the heterodimerisation of PSII during the early Archean.

The core of Type I reaction centres in anoxygenic phototrophic bacteria is homodimeric, made of a single subunit, known as PshA in phototrophic Firmicutes, and PscA in phototrophic Chlorobi and Acidobacteria (15). Photosystem I (PSI) in Cyanobacteria is heterodimeric, made of two homologous core subunits named PsaA and PsaB (Figure 1). It is thought that the heterodimerisation process was an evolutionary response to oxygen (16-18). More exactly, it is proposed that the asymmetric fine-tuning of the redox cofactors allows back electron transfer reactions to occur safely, minimising the formation of triplet chlorophyll states that can lead to the production of reactive oxygen species (17). It can thus be postulated that the gene duplication event that led to the heterodimerisation of PSI occurred after the evolution of water oxidation and was driven by the presence of oxygen.

**Figure 1.**
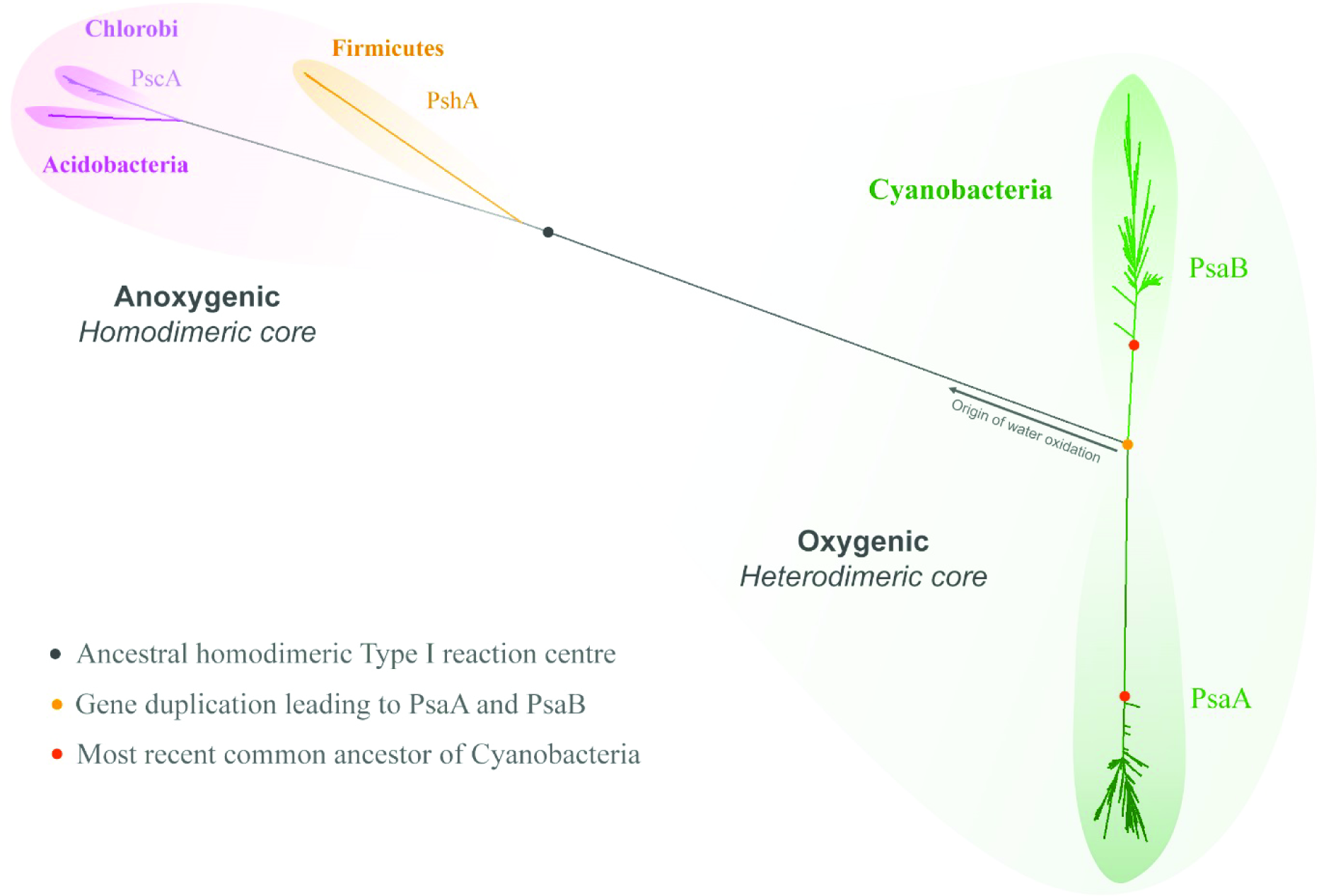
Maximum Likelihood tree of Type I reaction centre proteins. The tree is characterised by a deep split of reaction centre proteins, which separates those employed in anoxygenic phototrophic bacteria from those employed in oxygenic photosynthesis (grey spot). Cyanobacteria and photosynthetic eukaryotes are the only known phototrophs to have a heterodimeric Type I reaction centre made of two subunits known as PsaA and PsaB. All extant Cyanobacteria descended from a common ancestor that already had highly divergent PsaA and PsaB subunits (red spot). The gene duplication that led to PsaA and PsaB occurred at an earlier point in time (orange spot), which predated the most recent common ancestor of Cyanobacteria by an unknown period of time. It is hypothesised that the gene duplication that led to PsaA and PsaB occurred as a specialisation to oxygenic photosynthesis, therefore water oxidation should have originated before this gene duplication event (arrow).

In this report I present the first attempt at estimating when the gene duplication event that led to the heterodimerisation of the core of PSI occurred. I show that this gene duplication event is likely to have happened before 3.4 Ga, which is consistent with an early Archean origin of water oxidation.

## Results and discussion

### Sequence divergence of the core subunits of Photosystem I as a function of time

A first estimation of the time passed since the gene duplication event of the ancestral core subunit of PSI can be reached by comparing the level of sequence divergence between PsaA and PsaB in distantly related organisms. For example, in *Gloeobacter kilaueensis*, belonging to the earliest branching genus of Cyanobacteria capable of oxygenic photosynthesis (19, 20), PsaA and PsaB share 43.3% sequence identity (seq_id), see Table 1. In *Arabidopsis thaliana*, a flowering plant, PsaA and PsaB share 42.1% seq_id. In *Marchantia polymorpha*, a liverwort, and in *Cyanidioschyzon merolae*, a unicellular red alga, this value is 42.3%. The similarity of seq_id between PsaA and PsaB in these distantly related species suggests that in the very long time, likely more than 2.0 Ga ago, since the separation of the lineage leading to the genus *Gloeobacter* until the present time (3, 19), the rates of evolution of the core subunits have not varied drastically relative to each other, resulting in a difference of seq_id of only just over 1%.

**Table 1.**
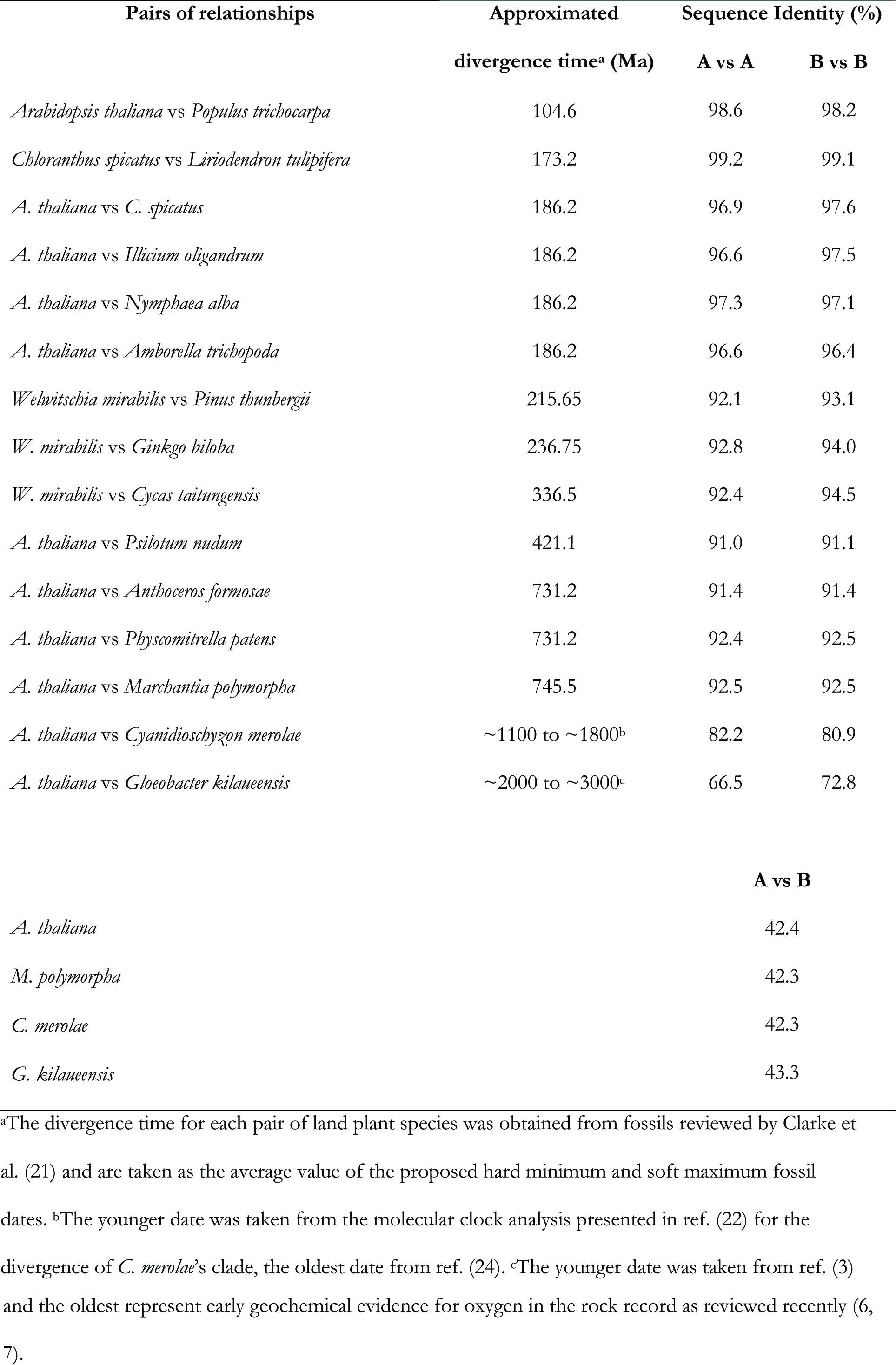
Percentage of sequence identity of PsaA and PsaB in various species

The PsaB subunit in *G. kilaueensis* and in *A. thaliana* share 72.8% seq_id and the PsaA subunit between these two species share 66.5% seq_id. Then, let us assume for the sake of simplicity that the most recent common ancestor (MRCA) of Cyanobacteria, at the divergence of the genus *Gloeobacter*, occurred around the time of the GOE: 2.4 Ga. This would mean that since the branching of *Gloeobacter*, at 2.4 Ga, PsaB subunits have only changed by a maximum of 27.2% and PsaA by about 33.5%. Again for simplicity, let us average this to 30%. It follows then that since 2.4 Ga the core subunits of PSI, PsaA and PsaB, have changed roughly about 30% each. That would give an average rate of evolution of about 1% loss of seq_id per every 80 Ma.

Using this rate we can then predict that two species that diverged about 100 Ma ago should have a level of sequence identity of their PsaA and PsaB subunits of just below **99%**. *A. thaliana* and *Populus trichocarpa* are known to have diverged about 100 Ma ago, or sometime between 82 and 127 Ma as inferred from fossil evidence (21). The PsaA seq_id between *A. thaliana* and *P. trichocarpa* is **98.6%** and that of PsaB is **98.2%**, which is in agreement with our approximated rate.

The oldest potential (though equivocal) fossils for crown group land plants (Embryophyta) are 1024 Ma and the oldest unequivocal fossil of a land plant is thought to be 449 Ma as discussed in detail by Clarke et al. (21). Thus it should be expected that the level of seq_id would be between **87.2%** based on the oldest fossil (1024/80 = **12.8%**) and **94.4%** on the youngest fossil evidence (449/80 = **5.6**%). The PsaA seq_id between *M. polymorpha*, an early branching land plant and *A. thaliana* is 92.5%. The same value is found for PsaB (Table 1). If the approximated rate is used to estimate the divergence of *M. polymorpha*, this would place the earliest land plants about 600 Ma ago. Molecular clock estimates for the divergence of *M. polymorpha* range from 410 to 680 Ma (21-24), which is in agreement with the approximated rate.

The earliest widely accepted fossil of red algae, the multicellular *Bangiomorpha pubescens*, was considered to be 1.2 Ga old (25). The date of this fossil was recently re-evaluated to 1.047 Ga (22). There is also a recently characterised putative multicellular red algae fossil 1.6 Ga old (26, 27). Using the approximated rate we can predict that the level of sequence divergence between *Cyanidioschyzon merolae* (a unicellular red algae) and *A. thaliana* should be below **86.9%** based on the younger fossil or about **80%** based on the older fossil (1047/80 = **13.1%** or 1600/80 = **20%**). In agreement with the approximated rate of evolution, the PsaA seq_id between these species is **82.2%** and the PsaB seq_id is **80.9%**. Using the approximated rate an age between 1.42 and 1.58 Ga for the divergence of the clade containing *C. merolae* can be extrapolated, which is similar to the age calculated by other molecular clocks (23) and to the age of the fossils mentioned above (26, 27).

From these comparisons it can be concluded that the rate of evolution of PsaA and PsaB has not changed dramatically for more than 2.4 Ga and that the approximated rate is not likely to be too far off from the average rates of evolution found in the wild.

Now, using this approximated rate we can extrapolate back in time to estimate when the gene duplication event that led to the divergence of PsaA and PsaB started. PsaA and PsaB share about 42.5% seq_id (Table 1), applying a rate of change of 1% loss of seq_id per 80 Ma gives a result of 4.6 Ga (57.5*80 = 4600 Ma). This is a very implausible date, but it is useful to illustrate a few key characteristics of the evolution of photosynthesis and reaction centre proteins as described in more detail below.

Let us assume instead that the MRCA of Cyanobacteria occurred at 3.0 Ga, 600 Ma before the GOE. That assumption would result in a slower rate of evolution of about 1% loss of seq_id for every 100 Ma. Consequently, it would set the gene duplication event at 5.75 Ga. In the other extreme case, if the MRCA postdate the GOE as suggested by Shih et al. (3), at 2.0 Ga, that would result in a faster rate of evolution of 1% loss of seq_id for every 66 million years. If this rate is applied to the divergence of PsaA and PsaB, it would set the gene duplication event at 3.79 Ga (57.5*66 = 3795 Ma), still long before the GOE. It becomes therefore apparent judging from a comparison of the level of seq_id alone that regardless of when the MRCA of Cyanobacteria existed (2.0 to 3.0 Ga), the gene duplication that led to PsaA and PsaB is more likely to be a tremendously ancient event, tracing back to the earliest stages in the evolutionary history of photosynthesis, than a late event occurring just before the GOE or after the GOE.

Figure 2 shows a plot comparing the level of seq_id of pair of species as listed in Table 1. It highlights that the rate of evolution of the core proteins of PSI have remained relatively stable for billions of years. From these basic considerations, and even allowing for very large uncertainties, it can also be concluded that any scenario for the divergence of PsaA and PsaB occurring less than 3.7 Ga ago would likely require faster rates of evolution at the earliest stages of divergence than any observed since the Proterozoic.

**Figure 2.**
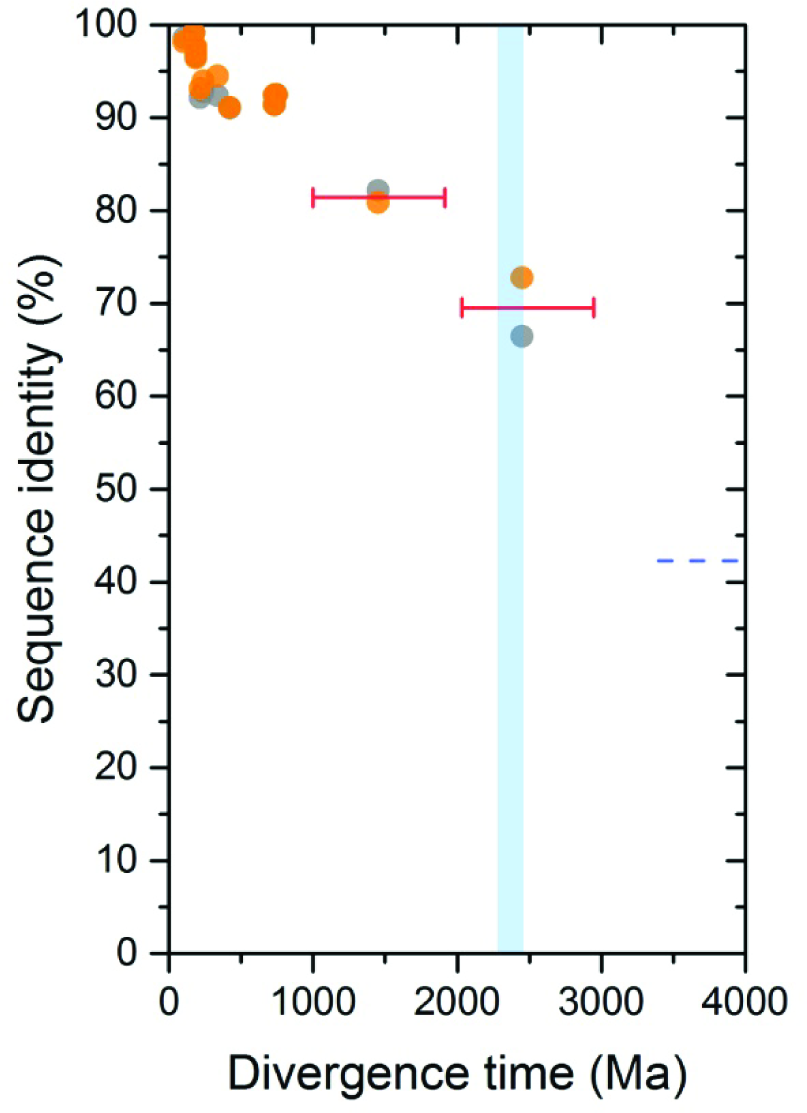
Percentage of sequence identity of PsaA and PsaB as a function of time. The data points are listed in Table 1. Dots in orange represent the level of sequence identity of PsaA from one species compared to the same subunit in a different species. Dots in grey represent PsaB for similar pairs of species. The blue bar represents the GOE. The oldest pair of dots are those from *G. kilaueensis* and are placed arbitrarily at about the start of the GOE. The second oldest node was set at 1.45 Ga and taken as the average of the estimated divergence time for the clade containing *Cyanidioschizon merolae* as calculated in ref. (22) and (24), dated in the range of 1.1 to 1.8 Ga respectively. The red bars represent the level of uncertainty on the estimated divergence times of the clades that contain these species. The blue dashed line marks the average level of seq_id between PsaA and PsaB in all known oxygenic phototrophs, at about 42.5%.

### Bayesian relaxed molecular clock analysis

To validate the above observations I performed a Bayesian molecular clock analysis on Type I reaction centre proteins assuming a root of either 3.5 ± 0.05 or 3.8 ± 0.05 Ga (Figure 3). In other words, assuming that photochemical reaction centres originated in the early Archean, which is consistent with extensive isotopic and sedimentological evidence for photoautotrophic metabolism during this time (2, 9, 10, 28-33). Calibration points are numbered and marked with red dots in Figure 3, are listed in Table 2, and described in Materials and Methods. The estimated divergence time of the duplication that led to PsaA and PsaB when constraining the MRCA of Cyanobacteria before the GOE was in both cases the oldest node after the root. An estimated divergence time of 3.41 ± 0.09 Ga was computed for the younger root and 3.62 ± 0.14 Ga for the older root (Table 3). The estimated divergence time for each node of each tree used in this analysis is supplied in Supplementary Data 1.

**Figure 3.**
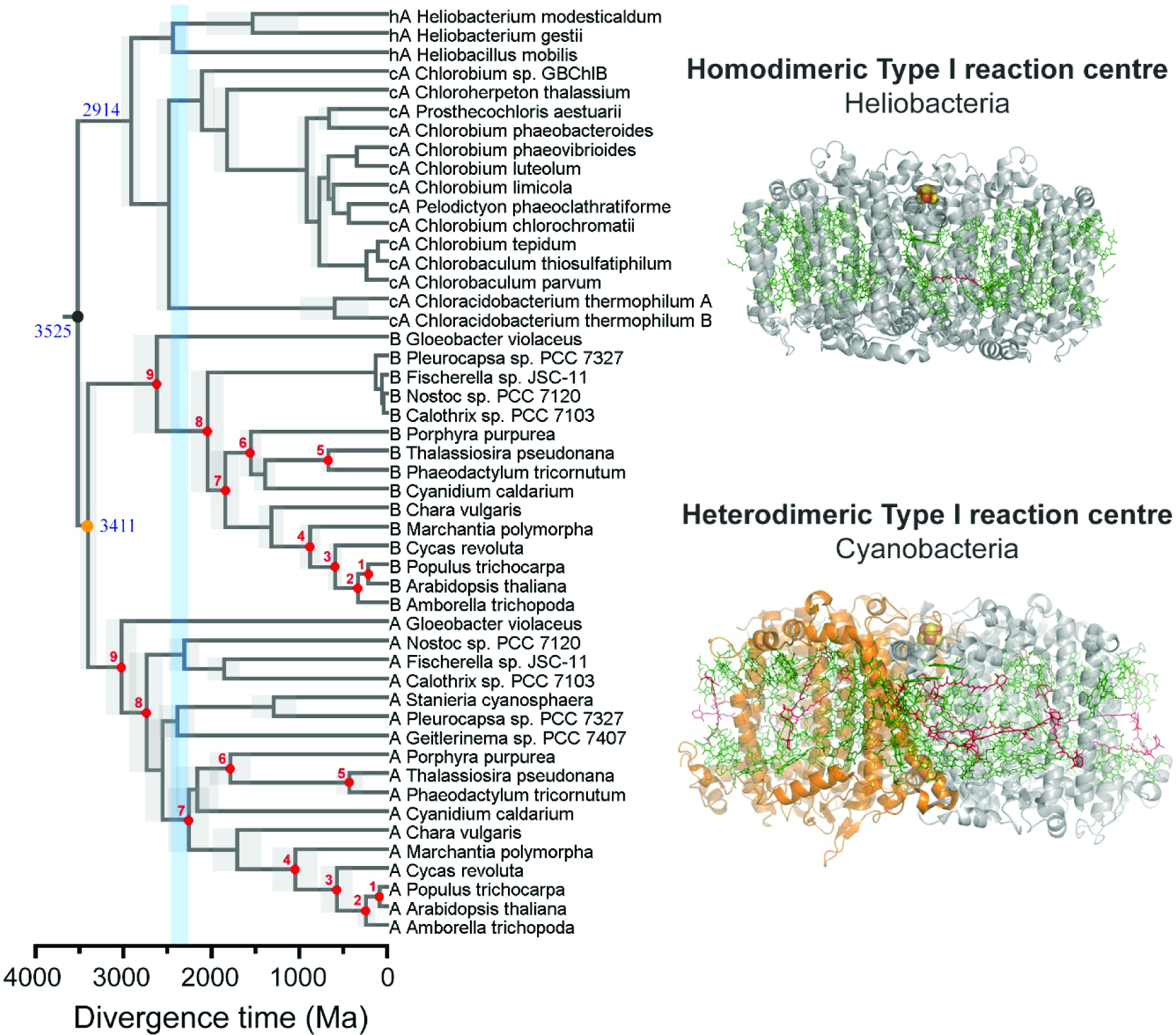
Bayesian relaxed molecular clock of Type I reaction centres. The tree was calculated assuming that Type I reaction centres had originated by 3.5 Ga (grey dot). The orange dot indicates the duplication event that allowed the divergence of PsaA and PsaB. Red dots highlight the nodes that were calibrated as described in Materials and Methods. The light grey bars along the nodes show the uncertainty, 95% confidence interval, on the estimated divergence time. Sequences marked as hA denote those from strains of Heliobacteria, while those marked as cA from Chlorobi and Acidobacteria. Sequences marked as A and B represent PsaA and PsaB from Cyanobacterial and eukaryotic PSI. The blue bar marks the GOE.

**Table 2.**
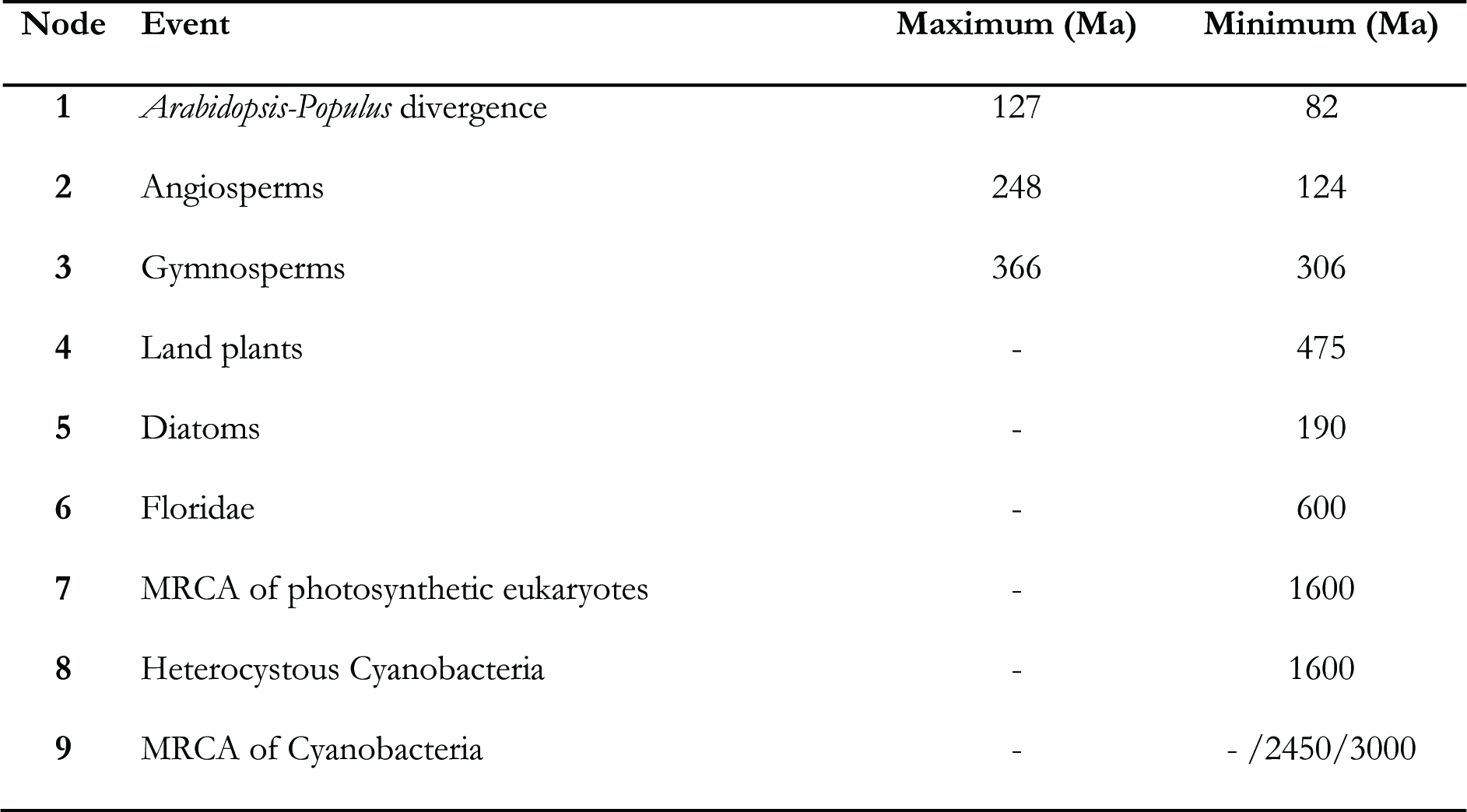
Calibration points

To test the effect of the calibration points (Figure 3, point 7, 8, and 9) on the estimated divergence time for the split of PsaA and PsaB, I repeated the same analysis this time constraining the oldest point to a minimum age of 3.0 Ga or alternatively removing the three oldest nodes altogether (Table 3). The plots shown in Figure 4 indicate that the calibration choice for the MRCA of Cyanobacteria had a small or no effect on the timing of the gene duplication event of the core subunits of PSI. Removing the three oldest node left no calibration older than 600 Ma on PsaA and PsaB, this setting has the effect of producing much younger ages for many nodes (Figure 4 C and F) and of increasing the level of uncertainty, yet the PsaA and PsaB divergence time still remained in the early Archean. In this case, a divergence time of 3.23 ± 0.21 and 3.34 ± 0.36 was estimated for the trees calculated using a root of 3.5 and 3.8 ± 0.05 Ga respectively.

**Figure 4.**
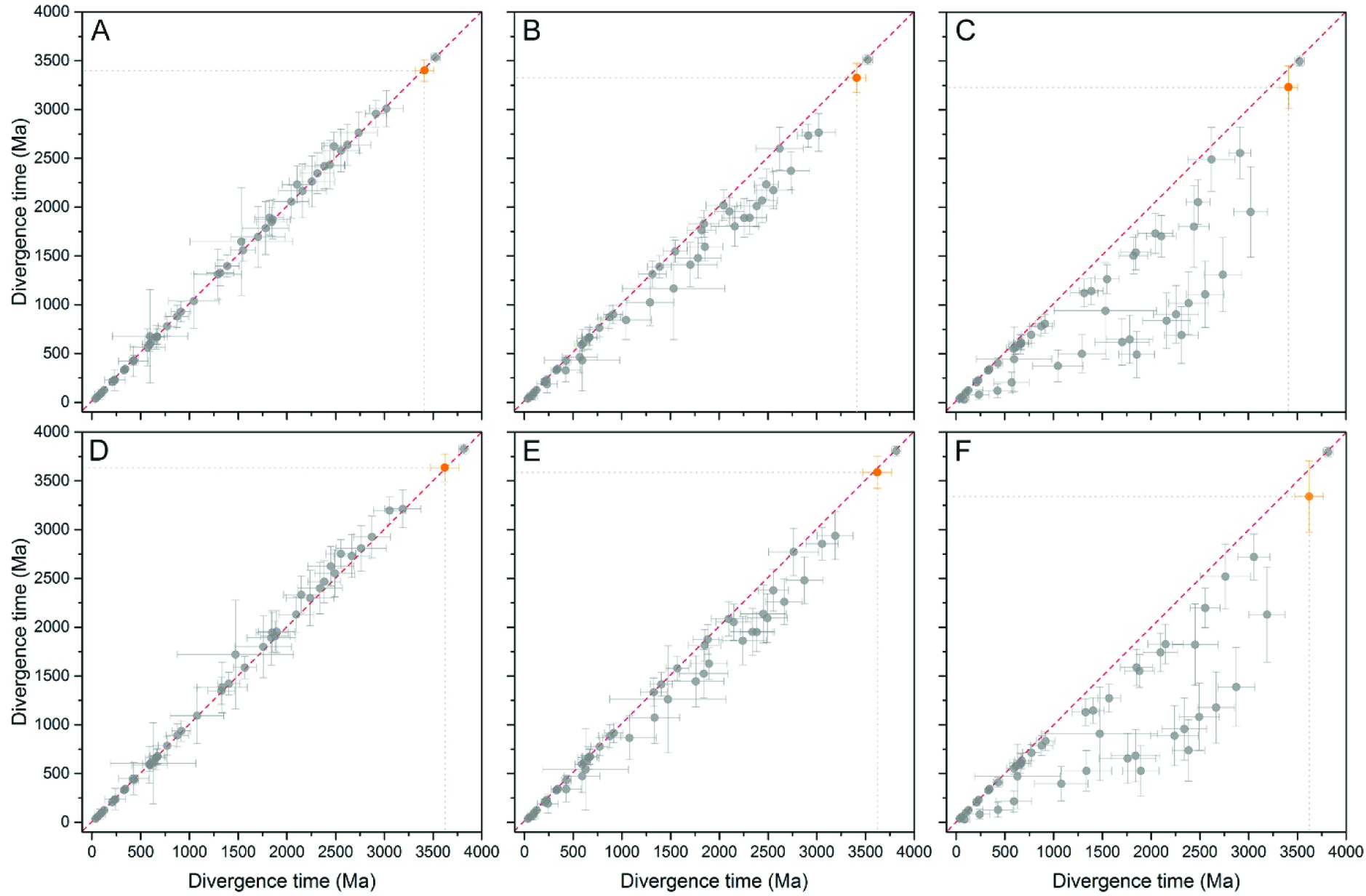
Effect of calibration choice on the estimated divergence time. (A) Comparison of divergence time estimates of a tree calculated using a root of 3.5 Ga and a minimum age for the MRCA of Cyanobacteria of 2.45 Ga (*x* axis) against a tree calculated using a minimum age for the MRCA of 3.00 Ga (*y* axis). There is no deviation from the slope (red dashed line) meaning that the calibration did not have a strong effect on the estimated divergence time. The orange dot represents the divergence time of the duplication that led to PsaA and PsaB. (B) Divergence times of a tree calculated using a root of 3.5 Ga and a minimum age for the MRCA of Cyanobacteria of 2.45 Ga (*x* axis) against a tree calculated without a calibration on this point (*y* axis). (C) Divergence times of a tree calculated using a root of 3.5 Ga and a minimum age for the MRCA of Cyanobacteria of 2.45 Ga (*x* axis) against a tree calculated without a calibration on points 8, 7, 9 (*y* axis). (D), (E), and (F) were calculated as (A), (B), and (C) respectively, but employing a root of 3.8 Ga instead. In all scenarios the duplication that led to PsaA and PsaB is the oldest node after the root. Removing the oldest calibration constraints led to younger ages on a number of nodes, however the estimated age of the PsaA/PsaB duplication event (orange dot) is set in the early Archean in every scenario.

**Table 3.**
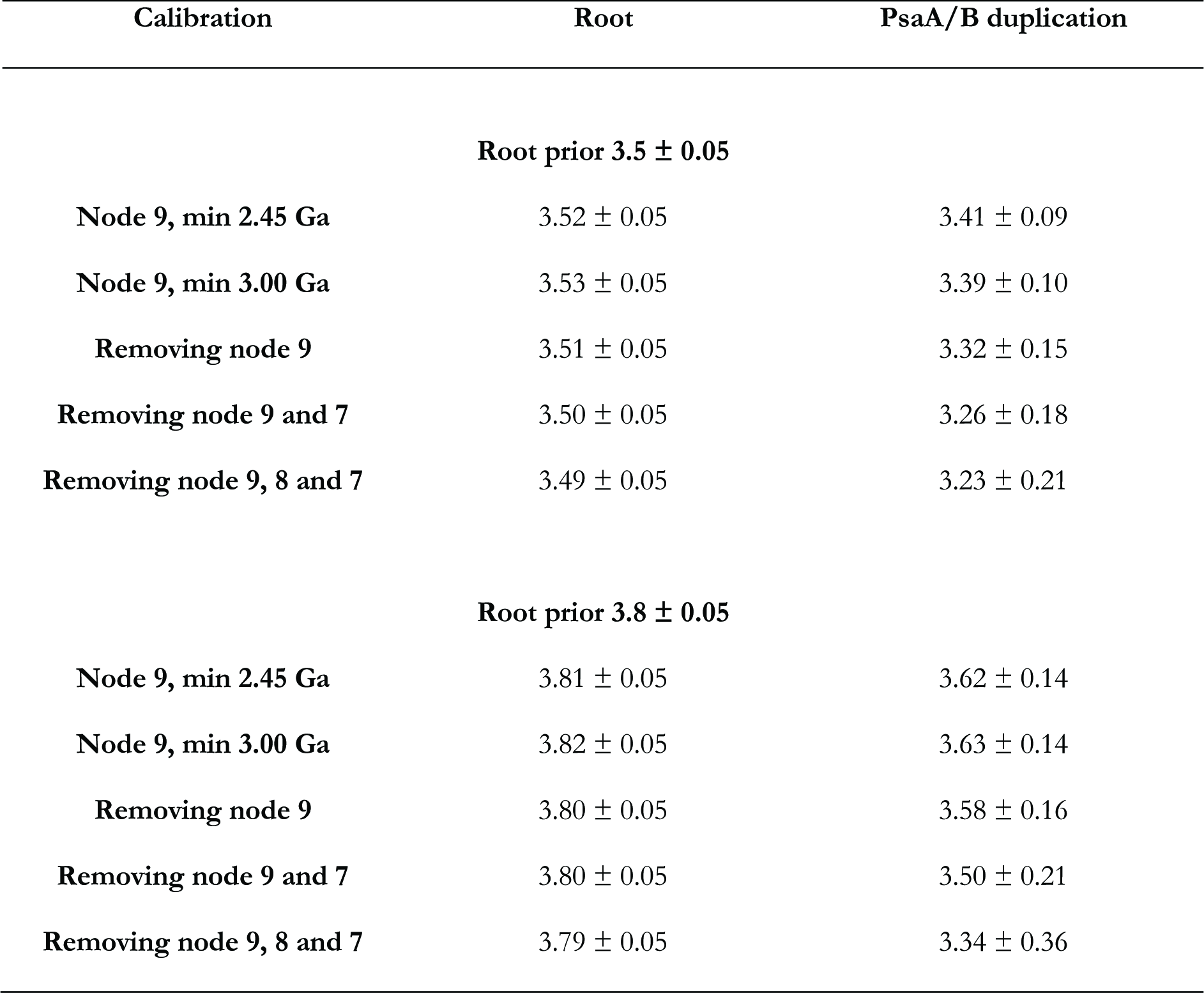
Effect of calibration points on the PsaA/PsaB duplication event

Type I reaction centres are characterised by having a light-harvesting domain (antenna) comprising the first six transmembrane helices and an electron transfer domain (core) comprising the last five transmembrane helices (34). To test whether the different selective pressures on the antenna or the electron transfer core could result in differences in the estimated divergence times, I performed molecular clocks using only the antenna or the core domain parts of the sequences (Supplementary Figure 1). No major differences were noted, and in each case the estimated age of the PsaA and PsaB divergence remained in the early Archean as the oldest node after the root.

Consistent with the conclusions derived from the comparison of the level of seq_id, even such ancient origins for the initial divergence of PsaA and PsaB still require high rates of evolution at the earliest stages. If the origin of photosynthesis happened about 3.5 Ga ago, that would require a rate at the time of the duplication event that led to PsaA and PsaB of 2.98 ± 0.79 amino acid changes per site per Ga, which is about 24 times larger than the average rate observed during the Proterozoic (0.12 ± 0.03 changes per site per Ga). If the origin of photosynthesis happened about 3.8 Ga ago, that would only slow down the initial rate to 2.76 ± 0.78 changes per site per Ga, with virtually no change observed for the average rate seen in the Proterozoic (Figure 5). This result implies that the phylogenetic distance between PsaA and PsaB is too large and the rate of evolution too slow, for the gene duplication event to have occurred within a few million years or a few hundred million years before the GOE. This is also in good agreement with the evolution of Type II reaction centres, which indicated that a water-splitting homodimeric photosystem is likely to have arisen in the early Archean (12, 13).

**Figure 5.**
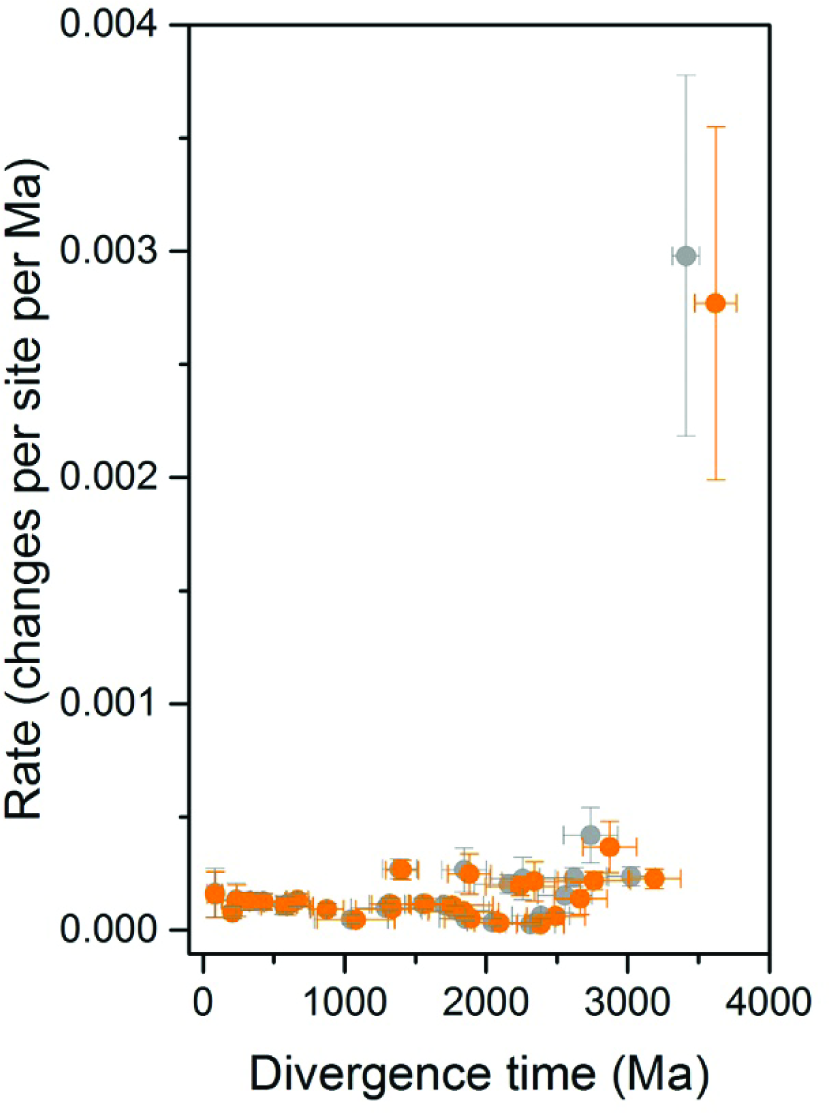
Rates of evolution as a function of divergence time. Grey dots represent divergence times of a tree calculated using a root constraint of 3.5 Ga (shown in Figure 3), while orange dots represent those calculated using a root constraint of 3.8 Ga. The fastest rate of evolution occurs at the PsaA/PsaB duplication event. The plot shows that higher rates of evolution, measured as amino acid changes per site per million years, are necessary to explain the divergence of PsaA and PsaB at any point in time in the history of life.

### Horizontal gene transfer

It has been suggested that Cyanobacteria could have obtained photosynthesis via horizontal gene transfer (HGT) from anoxygenic photosynthetic bacteria (35, 36). A recent version of this hypothesis suggests that a non-photosynthetic ancestor of Cyanobacteria acquired photosynthesis via massive HGT right before the GOE after the divergence of the Melainabacteria and the Sericytochromatia basal clades of non-photosynthetic Cyanobacteria (3, 37). The molecular evolution of Type I reaction centres is somewhat inconsistent with this hypothesis. This is because the ancestral PSI had to be a homodimer and as of today no anoxygenic bacteria with heterodimeric Type I reaction centres have been discovered, which suggests that the hypothesised HGT had to occur before the PsaA/PsaB duplication event. As we have seen above, it is unlikely that this duplication event occurred immediately before the GOE or after the GOE, given the fact that at this time PsaA and PsaB had already achieved a high degree of sequence divergence. These arguments also apply to Type II reaction centres as discussed in Cardona et al. (13).

Other possible evolutionary scenarios not completely ruled out by this analysis is that a very distant ancestor of Cyanobacteria obtained a homodimeric Type I reaction centre via HGT from a bacterium predating the divergence of PshA and PscA, long before the GOE. Another plausible scenario for HGT of PSI is that an ancestor of Cyanobacteria obtained a heterodimeric Type I reaction centre instead. This scenario would imply the existence of anoxygenic phototrophs (extinct or undiscovered) with heterodimeric Type I reaction centres and that the split of PsaA and PsaB was not related to the origin of photosynthetic water oxidation to oxygen. In such instance, an explanation for the initial divergence of PsaA and PsaB would need to be proposed, which does not invoke protection against the formation of reactive oxygen species.

Shih et al. (3) suggested based on molecular clock analysis that since the divergence of the Melainabacteria some 300 Ma years could have passed between the acquisition of reaction centres (from anoxygenic phototrophs) to the MRCA of crown group Cyanobacteria capable of oxygenic photosynthesis. Another recent molecular clock analysis indicated that this span of time could range from about 150 to about 560 Ma depending on the models and assumptions used (38). Both Shih et al. (3) and Magnabosco et al. (38) suggested that this could be enough time to develop oxygenic photosynthesis, but none of these studies analysed the evolution of the reaction centre proteins. While 300 or 560 Ma might seem like sufficient time, in terms of the evolution of reaction centre proteins it is not enough time. As shown in Table 1, 500 Ma is enough time to account for less than 10% sequence divergence between PsaA and PsaB. In the case of D1 and D2, the core of Photosystem II, 500 Ma is enough time to account for less than 5% sequence divergence (13). The MRCA of photosynthetic Cyanobacteria, defined by the divergence of the genus *Gloeobacter*, already had heterodimeric cores for both PSI and PSII, which had reached more than 55% sequence divergence for PsaA and PsaB, and more than 70% sequence divergence for D1 and D2. Therefore, not even one billion years before the MRCA of photosynthetic Cyanobacteria seems like sufficient time to account for such level of sequence divergence unless much faster rates of evolution are suggested at the earliest stages followed by an exponential decline in the rates. In fact, one billion years would still need rates of evolution many times greater at the earliest stages of diversification than any seen since the GOE to account for the large phylogenetic distance between PsaA and PsaB (this study), and between D1 and D2 (13). This evolutionary pattern can be inferred from a comparison of the levels of sequence identity alone and it is confirmed by Bayesian molecular clocks even allowing for large uncertainties on the calibration points. At this stage and to the best of my knowledge it is not known whether other proteins of very ancient origins show similar evolutionary patterns, but it was suggested recently that the spontaneous rate of mutation of early life could have declined exponentially as the planet cooled down (39), which would support an early origin of photosynthetic water oxidation, but other scenarios are possible.

### Heterodimeric Photosystem I and oxygen

When the electron acceptors of PSI are blocked under conditions of abiotic stress, back-reactions can occur leading to the formation of triplet chlorophylls, which can react with oxygen to produce singlet oxygen (17). Gisriel (40) suggested after the publication of the crystal structure of the homodimeric Type I reaction centre from the strictly anaerobe *Heliobacterium modesticaldum* (41), that every major structural change that occurred from the ancestral homodimeric reaction centre to cyanobacterial PSI can be explained by avoidance of singlet oxygen formation. Of particular note is a shift from loosely bound quinones in the ancestral homodimeric reaction centre (41) to entrapped and immobile quinones in PSI (42) to minimise the chance of charge recombination from the reduced acceptor chlorophylls (ec3^-^) back to P^+^. Because the phylloquinone binding site is highly conserved in PsaA and PsaB it was proposed that this modification for protection occurred at a homodimeric stage (40) (See Figure 6). This notion is also strengthen by the fact that PSI binds more carotenoids than the homodimeric reaction centre in close contact with numerous chlorophylls. For example, all six betacarotene molecules found within PsaA have identical positions in PsaB (Figure 6) suggesting that prior to the PsaA and PsaB duplication event, the homodimeric PSI had already acquired a greater capacity to quench chlorophyll triplets (42). In comparison, the heliobacterial reaction centre binds a single carotenoid per monomer (41). This is consistent with the evolution of water oxidation occurring before the PsaA/PsaB split.

**Figure 6.**
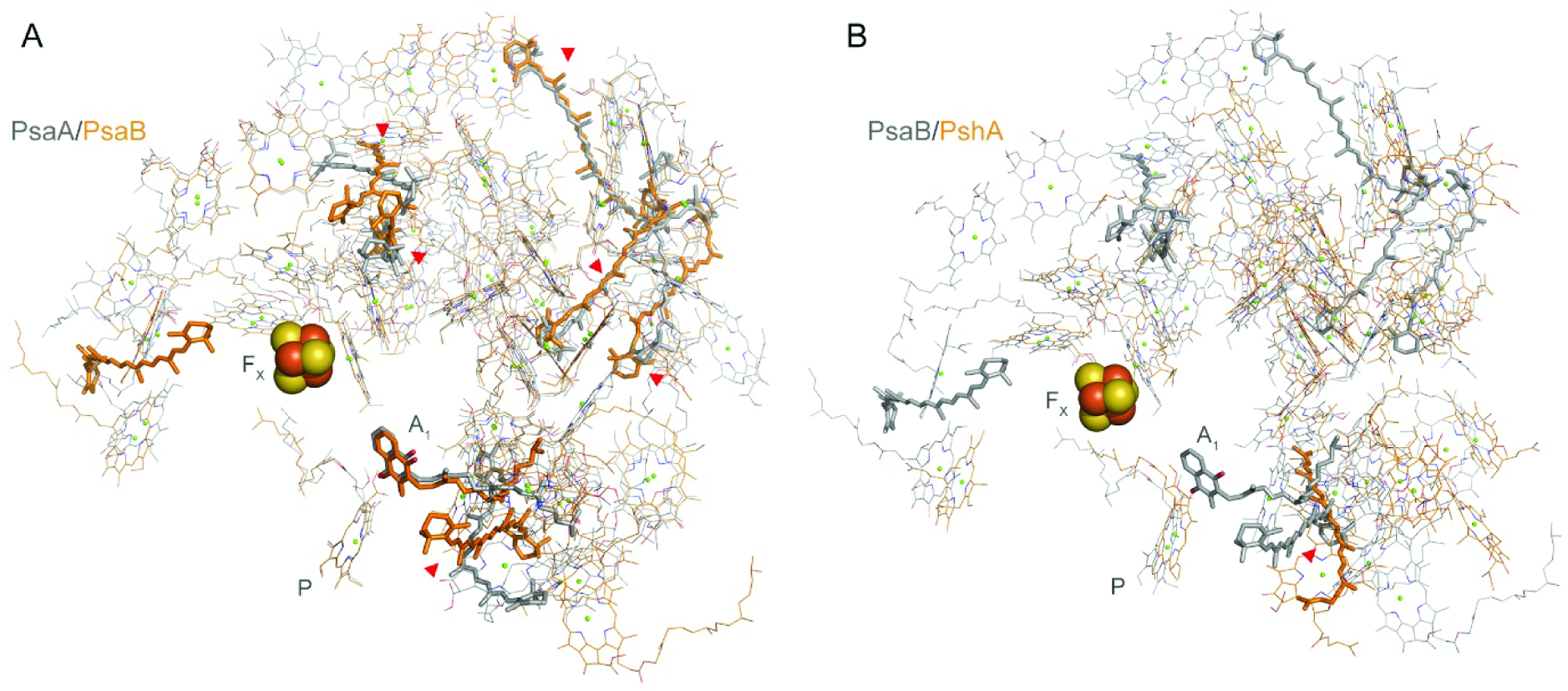
Structural comparisons of cofactor positions in Photosystem I and a homodimeric Type I reaction centre. (A) Overlap of PsaA (grey lines and sticks) and PsaB (orange lines and stick). Conserved carotenoids between PsaA and PsaB are highlighted with red arrows. Betacarotene molecules are shown in thick sticks and chlorophylls in thin lines. The phylloquinone cofactor is marked with A_1_, the Fe_4_S_4_ cluster is marked with F_X_, and the redox chlorophyll that makes the ‘special pair’ with P. (B) Overlap of PsaB and PshA from the anoxygenic phototrophic anaerobe *Heliobacterium modesticaldum.* The homodimeric Type I reaction centre lacks a quinone binding-site, has bacteriochlorophyll *g* as the main pigment, and features a single carotenoid molecule per monomer (4,4'-diaponeurosporene). The PDB ID for the structure of PSI is 1jb0 (42) and for the structure of the heliobacterial reaction centre is 5v8k (41).

In addition, Rutherford et al. (17) suggested that the difference in the reduction potential of the phylloquinones, estimated to be -531 mV for A_1A_ and -686 mV for A_1B_ (43), is an adaptation to allow the back-reactions to occur dominantly on the A branch from the semiphylloquinone radical directly to P^+^, thereby minimising the formation of chlorophyll triplet state and singlet oxygen. In contrast, a back-reaction via the B branch would result in chlorophyll triplet formation. The difference in the potentials is thought to be due to the asymmetry of the protein backbone and the presence of distinct ionisable amino acids in the vicinity of the phylloquinones (17, 43). Furthermore, the evolution of a heterodimeric PSI has also been attributed to the asymmetric binding of additional protein subunits at the onset of oxygenic photosynthesis (16), most importantly to optimise the binding of PsaC, which holds the two terminal Fe_4_S_4_ cluster (F_A_ and F_B_) and known to have evolved from a ferredoxin-like protein (44, 45). It is thought that PsaC and its strong binding to the PsaA/PsaB heterodimer evolved partly to shield and protect the Fe_4_S_4_ clusters against oxygen (45). Along the same lines, Gisriel (40) suggested that the permanent binding of PsaC, which is absent in the heliobacterial reaction centre, evolved to further separate the negative charge from P^+^ after charge separation. This would decrease the chance of back-reactions by preventing the accumulation of the semiquinone radical in the presence of reduced F_X_ under conditions that would lead to an over-reduction of the electron transport chain.

Taken together the above considerations for the evolution of a heterodimeric Type I reaction centre, it seems quite plausible that the gene duplication event that allowed the divergence of PsaA and PsaB occurred only after the emergence of water oxidation to oxygen in an ancestral Type II reaction centre. What is more, it would appear as if the ancestral Type I reaction centre of oxygenic photosynthesis had acquired some protection mechanisms still in a homodimeric state.

### Final remarks

The high level of sequence divergence between the two core subunits of each photosystem, a trait shared by all Cyanobacteria, highlights the fact that their *most recent common ancestor* existed at an advanced stage in the evolution of oxygenic photosynthesis. This makes evident that a great diversity of oxygenic phototrophic bacteria spanning from the origin of water oxidation (using homodimeric reaction centres) to the emergence of crown group Cyanobacteria (using very sophisticated multiprotein photosystems with very divergent heterodimeric cores), either still remain undiscovered or have gone completely extinct. From the low level of sequence identity of the core subunits in addition to relatively slow rates of evolution it can be deduced that most of the diversity of oxygenic phototrophic bacteria actually predated crown group Cyanobacteria. Indeed, the evolutionary distance between the core subunits is so large that the early diversity of oxygenic phototrophic bacteria was likely not contained within a single ancestral phylum. Consequently, the possibility that Cyanobacteria are not the only existing phylum of bacteria to have descended from oxygenic phototrophs should be carefully considered in scenarios for the early evolution of bioenergetic processes and the interpretation of microfossils, stromatolites, and the geochemical record of early Archean rocks.

In conclusion, the data presented here indicates that the gene duplication event that allowed the heterodimerisation of Photosystem I—in an ancestral lineage of bacteria that predated the most recent common ancestor of the phylum Cyanobacteria—likely occurred in the early Archean. If such event was an adaptation to photosynthetic oxygen evolution it would set water oxidation more than a billion years before the Great Oxidation Event.

## Materials and Methods

All sequences used in this study were retrieved from the NCBI refseq database and are provided in the supplementary files titled: Supplementary Sequences 1 and 2. All sequence alignments were performed using Clustal Omega employing 10 combined guide trees and Hidden Markov Model iterations (46). The percentage of sequence identity between pair of sequences was estimated using the Sequence Manipulation Suite online service (47). The Maximum Likelihood tree using all retrieved sequences and shown in Figure 1 were calculated with PhyML 3.0 using the smart model selection option allowing the software to compute all parameters from the data based on the Bayesian Information Criterion (48). The topology of this tree is identical to other phylogenetic studies on Type I reaction centres and it has been reviewed extensively before by myself and others, see for example (14, 36).

A total of 59 sequences spanning the entire diversity of reaction centre proteins, including 10 PsaA and 10 PsaB sequences from photosynthetic eukaryotes, were selected for molecular clock analysis. These sequences include the earliest branches in the tree of Cyanobacteria, those of *Gloeobacter*, heterocystous cyanobacteria, and sequences from photosynthetic eukaryotes that are amenable for setting fossil calibrations as described in more detail below. Only very few sequences for Heliobacteria, Chlorobi, and Acidobacteria are publically available and all of these were included in the molecular clock analysis, with the exception of the PscA sequence from *Chlorobium ferrooxidans*, which appears to be incomplete at the C-terminus or poorly sequenced. The PscA sequence of *Chloracidobacterium thermophilum* is characterised by a sequence insertion between the 7^th^ and 8^th^ transmembrane helix of about 165 amino acids (49). This insertion was removed from the sequence alignment.

Calibration points were assigned as shown in Figure 3 and age boundaries were allocated as listed in Table 2. A minimum and a maximum divergence times for the *Arabidopsis*/*Populus* divergence, the appearance of the angiosperms (*Amborella* divergence), gymnosperms (*Cycas* divergene), and earliest land plants (*Marchantia* divergence) were obtained from Clarke et al. (21), which described the fossil record of land plants in great detail and suggested the best recommended practice. In the case of the earliest land plants, Clarke et al. suggested a maximum age of 1024 Ma, this calibration was not used and no maximum age was set for this node. A minimum age for diatoms was obtained from Jurassic fossils from the Lias Group, reviewed by Sims et al. (50). No maximum age was used for this node. Only a minimum age of 600 Ma was assigned to the node between the diatom sequences and the sequences from *Porphyra purpurea*. These were selected based on a Late Neoproterozoic Chinese multicellular red alga fossil (51) and I used it as a conservative estimate for the divergence of complex red algae, which are probably older (23). The well-preserved 1.6 Ga red algae fossils recently reported by Bengtson et al. (26, 27) was used as a minimum age for the origin of photosynthetic eukaryotes, an age which is in agreement with recent molecular clock analysis (23, 24). Although, this date was recently contested by Gibson et al. (22) who proposed an earlier origin of photosynthetic eukaryotes at about 1.2 Ga. In addition, a minimum age of 1.6 Ga was assigned to the divergence of heterocystous cyanobacteria as well-preserved akinetes of this age have been found in cherts from Siberia, China, and Australia (52-54). A minimum age of 2.45 Ga was assigned to the node at the divergence of *Gloeobacter* assuming that the most recent common ancestor of Cyanobacteria is at least as old as the GOE (55).

Currently, it is not known exactly when the MRCA of Cyanobacteria and photosynthetic eukaryotes originated. To consider a range of possibilities, I also performed molecular clocks using an older minimum age for the MRCA of Cyanobacteria of 3.0 Ga. This was done to test the possibility that some of the earliest evidence for oxygen (7) may have arisen from crown group Cyanobacteria. In addition, I also performed the same analysis removing all calibration points older than 1.0 Ga to release those nodes from any time bias.

When the earliest reaction centres originated for the first time also remains unknown. There is however extensive evidence for photoautotrophy about 3.5 Ga, see for example (8, 9, 33); yet it is plausible that the geochemical record of ancestral forms of photosynthesis extends much further back in time to about 3.8 Ga (2, 10, 28-32). I have therefore constrained the root of the tree to 3.5 or 3.8 Ga to compare the effect of the root priors on the estimated divergence times.

A Bayesian Markov chain Monte Carlo approach was used to infer estimated divergence times with the Phylobayes 3.3f software (56). A relaxed log-normal autocorrelated molecular clock was applied together with the CAT model of amino acid change and applying a uniform model of amino acid substitutions (Poisson) (56, 57). The relaxed autocorrelated clock model allows the rate of evolution to vary from branch to branch but assumes that closer branches are more likely to evolve at similar rates (58). The CAT model can account for substitutional rate heterogeneity across the sites of the protein sequences (56, 57). Soft bounds on the calibration points were used for maximum flexibility on the time constrains and were left as default; that is 2.5% tail probability falling outside each calibration boundary or 5% in the case of a single minimum boundary (59). All molecular clocks were computed using four discrete rate categories for the gamma distribution and four chains were run in parallel until convergence. Rates of evolution, expressed as amino acid changes per site per unit of time, are inferred by the software as described by the developers elsewhere (60, 61).

